# DiffMethylTools: a toolbox of the detection, annotation and visualization of differential DNA methylation

**DOI:** 10.1101/2025.07.01.662655

**Authors:** Houssemeddine Derbel, Evan Kinnear, Justin J.-L. Wong, Qian Liu

## Abstract

DNA methylation is a fundamental epigenetic mechanism, and its significant changes (i.e., differential methylation) regulate gene expression, cell-type specification and disease progression without altering the underlying DNA sequence. Differential methylation was usually detected via existing statistical tools by comparing two groups of methyomes (i.e. whole-genome methylation profiles) and has wide applications of various downstream investigations for human disease studies. However, few toolboxes were available to efficiently streamline methylation investigation by integrating robust detection, annotation and visualization of differential methylation. Also, differential methylation detected via tools has poor reproducibility and no tools were tested on the increasing volume of long read methylomes. To address these issues, we introduced DiffMethylTools, an end-to-end solution to eliminate analytical and computational difficulties for differential methylation dissection. Comparison of detection performance on six datasets including three long-read methylomes demonstrated that DiffMethylTools achieved overall better performance of detecting differential methylation than existing tools like MethylKit, DSS, MethylSig, and bsseq. Besides, DiffMethylTools supported versatile input formats for seamless transition from upstream methylation detection tools, and offered diverse annotations and visualizations to facilitate downstream investigations. DiffMethylTools therefore offered a robust, interpretable, and user-friendly solution for differential methylation investigation, benefiting the dissection of methylation’s roles in human disease studies.

## 1. Introduction

DNA methylation is a compulsory and fundamental epigenetic process and regulates gene expression and cellular function without altering the underlying sequence ^1^. DNA methylation, especially 5-Methylcytosine (5mC), widely occurs in mammal genomes ^2^. 5mC is recognized as the fifth type of nucleotides besides the four standard nucleotide types in the human genome, and presents at most twenty-nine million CpG sites. DNA methylation thus plays crucial roles in epigenetic regulation and is essential for biological processes such as genomic imprinting^3^, X-chromosome inactivation^4^, and suppression of transposable elements^5^. DNA methylation also stably regulates cell type and tissue specific biological functions of an identical genome in an individual^6^, highlighting the importance of DNA methylation in biological systems.

Whole genome profile of DNA methylation could be detected via different sequencing techniques, such as widely-used bisulfite sequencing (BS-seq) and reduced representation bisulfite sequencing^7^ (RRBS), which used bisulfite conversion and short-read sequencing to detect methylated and unmethylated sites. Recently, third-generation sequencing (3GS) technologies, including Nanoball Sequencing ^8^ (Illumina’s Nanoball Technology), Helicos Sequencing^9^ (Helicos BioSciences), Single-Molecule Real-Time Sequencing^10^ (SMRT) from Pacific Biosciences, and Oxford Nanopore Technologies ^11,12^ (ONT), were proposed to improve the methylation detection especially in low-complexity genomic regions and to enable the detection of non-5mC DNA modifications.

The advancement of methylation detection stimulated the detection of differential methylation patterns across various biological process and human diseases^3,13–15^. These differential methylation patterns offered valuable insights into gene expression regulation influenced by environmental factors^16^, developmental processes^17^, or disease states^18^, and facilitating the discovery of biomarkers for early disease diagnosis^19^, prognosis^20^, and monitoring of disease progression^21^ or treatment response^22,23^.

There are two types of differential methylation patterns, differentially methylated loci (DMLs) for differential CpG sites or positions, and differentially methylated regions (DMRs) which are a cluster of DMLs and reflect synergistic effect of DMLs. These methylation patterns could be detected by various computational tools using whole-genome methylation profiles as input. These tools could be classified into five categories according to the techniques used. One category of tools used simple statistical models like Fisher’s exact test and t-tests to compare methylation frequencies between conditions. These methods, such as DMRfinder^24^, and eDMR^25^, were fast and worked well with high coverage data. But these models ignored biological variability. The second category of computational tools used generalized linear model (GLM)-based methods, and included methylKit^26^, methylSig^27^, DSS-single^28^, DSS^29,30^, DMRcaller^31^, MOABS^32^, and RADMeth^33^. These methods modeled methylation counts or methylation levels using binomial or beta-binomial distributions to account for both biological variability and technical noise. These methods were generally well-suited for high-coverage sequencing datasets, and their performance tended to degrade on low-coverage datasets where variance estimates become unreliable. The third category of computational tools, such as DMAP^34^, RnBeads^35,36^ and COHCAP^37^, used linear regression to model methylation levels across conditions with an assumption of normal distribution of methylation. They worked well on high-quality, high-coverage data, such as array-based platforms or deeply sequenced bisulfite data. The fourth category of computational tools, such as such as BSmooth^38^ and BiSeq^39^, used smoothing process to reduce random noise by aggregating nearby CpG sites and were effective on low coverage data. However, smoothing did not show much improvement compared to other tools^40^. The fifth category of computational tools used Hidden Markov Model^41^ (HMM) (such as HMM-DM^42^, HMM-Fisher^43^ and Bisulfighter^44^) to model methylation states and transitions along the genome. These techniques worked well with short-range dependency and could handle low-coverage data better than binomial-based methods. However, these techniques are computationally intensive and difficult to generalize. In summary, diverse assumptions in different methods led to poor overlap of detected differential methylation patterns. Besides, most of these tools either lack built-in modules or offer limited support for annotation and visualization. For example, many tools could not conduct annotation or visualization for differential methylation, although methylKit relied on an external package for simple annotation tasks. A few packages, such as DMRcaller, MethylSig, and COHCAP, offered downstream processes of plotting local methylation profile, but they typically provided either basic annotation or rudimentary visualization. That is, most tools remained focused exclusively on differential methylation detection, leaving users to assemble separate, often fragmented pipelines for biological interpretation and visualization. Thus, a toolbox is needed to simplify and speedup the detection, annotation and visualization of differential methylation patterns.

To address these issues, we designed DiffMethylTools, a single-command-for-all tool for differential analysis. DiffMethylTools is flexible to various input formats of DNA methylation and performs reliable detection of DMLs and DMRs as well as annotation and visualization in a single command. We evaluated DiffMethylTools on two WGBS short-read data and three long-read sequencing data generated in our lab, and compared its performance against four widely used tools---MethylKit, DSS, MethylSig, and bsseq. The results demonstrated that DiffMethylTools overall outperformed the existing tools to detect benchmark methylation patterns. DiffMethylTools thus offered straightforward investigation of differential methylation patterns and will benefit human disease studies to investigate epigenetic roles in disease development and progression. DiffMethylTools is publicly available via https://github.com/qgenlab/DiffMethylTools.

## 2. Methods and Datasets

### 2.1 Datasets

#### 2.1.1 Whole-genome methylation data

To evaluate the performance of DiffMethylTools and existing tools in detecting differential methylation patterns, we collected five diverse datasets generated by ONT long-read sequencing or bisulfite sequencing short-read sequencing: One ONT data was for a HCT116 cell lines and its knockout of inositol polyphosphate multikinase (IPMK) gene^45^. This dataset included two biological replicates per condition (HCT116 wide type and knockout). Prior studies have shown that IPMK depletion leads to widespread alterations in DNA methylation^45^, making it suitable for evaluating the tool’s sensitivity to detect abundant methylation changes. The second ONT dataset was for non-differentiated THP-1 monocytes (called MO) and their differentiation of macrophage-like cells (ND). Each cell type contains three sequencing replicates. The third long-read dataset was for 12 patients with Alzheimer’s disease (AD) and eight unaffected controls. These three datasets were generated by us and the last two were not published. The rest two methylation datasets were generated by short-read bisulfite sequencing technologies. One is the published liver cancer dataset with four liver tumor samples and four matched control samples^46^. and the other was for cell-type samples of the Natural Killer (NK) versus B-cell, each containing three replicates^47^. The sequencing depth information of these datasets was summarized in Supplementary Figures 1-5. Together, these datasets provided a comprehensive benchmark for evaluating the effectiveness, generalizability and flexibility of detection tools across a range of different sequencing platforms, experimental conditions and biological contexts as well as various formats of methylation profiling.

Preprocessing of ONT data: After ONT raw data were generated using P2-solo and basecalled via ONT’s Dorado basecaller (version 7.2.13+fba8e8925), we used ONT’s modbam2bed to summarize the whole-genome methylation profiling and then computed coverage and methylation percentages using the equations provided below.

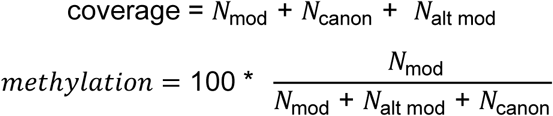

Where *N*_mod_, *N*_canno_ and *N*_alt_mod_ were count of reads with and without methylation calling as well as with an alternative methylation calling, respectively.

#### 2.1.2 Simulation data

We also generated simulation data with known DMLs to evaluate the performance of differential methylation detection tools. To preserve key biological characteristics observed in real methylomes, we used one methylation data from the HCT116 dataset as a starting methylome to simulate case and control profiles. First, we clustered CpG sites into regions, allowing a maximum gap of 100 base pairs between adjacent sites, and retained only regions containing at least 20 CpG sites. Within each region, methylation levels were smoothed using the LOWESS (locally weighted scatterplot smoothing)^48^ algorithm to obtain baseline methylation profiles.

To simulate control methylomes, methylation levels at individual CpG sites were sampled from a truncated normal distribution. This normal distribution had a mean of the local smoothed methylation level and a standard deviation (σ) that was independently generated from a uniform distribution ranging from 2 to 15. Directionality of methylation (hyper- or hypo-methylation) was assigned at random.

To simulate case methylation data, we introduced diverse methylation changes across the genome. Specifically, we assumed that 40% of CpG sites had minimal change between case and control conditions, 30% had ±5% difference, 10% by ±10% difference, and another 10% by ±15% difference. The remaining sites had larger methylation differences of ±20%, ±30%, ±40%, and ±50%. These case simulation samples were generated via the similar process for the simulation control samples: methylation percentages for each CpG site were drawn from a truncated normal distribution with the mean equal to the smoothed methylation level ± methylation difference and the standard deviation σ randomly selected from a uniform distribution between 2 and 15, and a random assignment was selected for the direction (±) of methylation change (hyper- or hypo-methylation). We generated three biological replicates for the case group.

To mimic real methylation data, we also simulated coverages for each CpG site. The simulation coverage for a CpG site was calculated via the equation below using an exponential decay function with Gaussian noise based on the absolute difference between the real and simulated methylation levels.

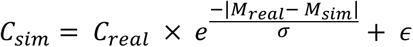

Where *C*_*sim*_ was a simulated coverage, *C*_*real*_ was a real coverage from the simulation starting methylome, σ represented the base standard deviation of the methylation at a given position, *ε* was a Gaussian noise, *M*_*real*_ was the methylation level from the smoothed reference profile in the control dataset or the smoothed methylation levels after methylation changes in the case dataset, and *M*_*sim*_ was the simulated methylation level for that position within the respective dataset. Using this formula, we assumed that the coverage of a CpG site was higher if its methylation level closely matches the average smoothed level. Overall, our simulation strategy yielded methylation and coverage profiles that mimic real sequencing data, providing a robust foundation for benchmarking differential methylation tools.

### 2.2 Flowchart of DiffMethylTools

DiffMethylTools was comprised of several sequential steps, including methylation preprocessing with noise filtering, DML detection, DMR detection, differential methylation annotation as well as differential methylation visualization, as described below.

#### Methylation preprocessing

The inputs of our DiffMethylTools were two groups of methylation profiles: one group for case samples and the other group for control samples. Each methylation sample was organized in a BED format of CpG sites together with methylation coverage, total coverage and/or methylation percentages. In DiffMethylTools, users could provide different formats to define methylation levels and their coverage. For example, a BED format could contain methylation coverage and total coverage, or methylation percentages and total coverage in various orders and different fields. This flexibility was useful because preprocessing tools, like Bismark and modbam2bed, usually did not use the same output format, which needed annoying but necessary data preprocessing before differential methylation detection.

The sites from each input methylome were first filtered with the coverage threshold of *c*1.

That is, any sites with less coverage were filtered out. Then, the input methylomes were combined to a matrix 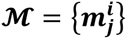 where each row represented a site ***j***, and a column represented a methylome sample ***i*** (***0*** ≤ ***i*** < ***N***), and 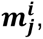 ranging from 0 to 100, was the methylation percentage (*i.e.*, the ratio of methylation calling over the total coverage for site ***j*** in a sample ***i***), and ***N*** was the number of input samples of two groups. After that, we used another coverage threshold *c*2 ≥ *c*1 to filter out those sites which had less than *t* methylomes with the coverage ≥ *c*2 from either the case or control group. The two groups could have different thresholds of *t* if the numbers of methylomes in the two groups were different.

In human genomes, there were more than 29 million of CpG methylation sites and majority of them had small methylation variability across and/or within the groups of samples ^49^. To focus on DML candidates and reduce running time, we filtered out those sites if their standard deviations of all samples in ***M*** were smaller than δ (default 10, adjustable). Standard deviations were calculated using methylome samples from both groups. After that, the missing values were filled with the group means.

#### DML detection

We fed ***M*** to a linear regression model implemented in Limma ^50^. In Limma, empirical Bayesian was used to fit the methylation using groups and other co-factors such as ages if available. A statistical p-value was calculated for each site to determine a significant association of methylation differences and the groups. These p-values were corrected using multiple testing strategy such as the Benjamini-Hochberg^51^ procedure to control the False Discovery Rate (FDR). A site with corrected p-value<=0.05 was considered as DMLs.

#### DMR detection

Many CpG sites remain proximal in DNA sequences and existing studies found the co-methylation^52–55^ of adjacent CpG sites, suggesting co-localization effect. We thus clustered DMLs into DMRs using the criteria below: (1) a 1000bp region contained more than a minimum number of DMLs (default: 3) and their direction of methylation changes was consistent (either all hypomethylated or all hypermethylated), (2) <30% methylation sites within the region exhibited random methylation changes between two groups (the absolute difference of group means <7.5 by default), (3) <10% methylation sites in the region had opposite methylation changes (>=7.5 methylation levels) compared to the DMLs in this region. That is, if the DMLs in the region were hypomethylated, hypermethylation change was the opposite change; otherwise, hypomethylation change was the opposite change. Any regions that did not meet these criteria were split into smaller regions for similar clustering processing. Resultant regions were called DMRs.

#### Methylation annotations

To facilitate the investigation of biological impact of differential methylation, the detected DMLs and DMRs were annotated with known biological genomic regions using gene annotations from GENCODE (release 42), candidate cis-Regulatory Elements (cCREs) from ENCODE data^56^, repeat categories provided by UCSC genome browser^57^, and CpG sites from Illumina’s Infinium MethylationChip (HM450, EPIC v1 and v2). These annotations were downloaded and anchored to all 29,401,795 CpG sites in hg38. Then, these annotations were linked to DMLs and DMRs as described below.

The link of DMLs/DMRs to MethylationChip enabled easy validation of differential methylation in epigenome-wide association studies (EWAS) if available. Pie charts were used to show how many DMLs/DMRs could be detectable in EWAS, given that EWAS usually detected <3% of all CpG sites. Annotations of repeats to DMLs/DMRs were useful to investigate potential methylation changes in repeat regions.

Gene annotations of DMLs and DMRs included two components. First, we overlapped different methylation patterns with gene definitions, such as UTR (untranslated region), CDS, introns, exons, upstream regions and intergenic regions. If a methylation pattern overlapped with non-intergenic regions, this pattern was linked with this gene. A pattern could overlap with multiple genes. Second, we overlapped methylation patterns with enhancers and promoters. Promoter methylation typically silences genes by blocking transcription factor binding, while enhancer methylation disrupts long-range activation. After enhancers or promoters were anchored with a methylation pattern, a gene was annotated with the methylation pattern if it was in 2kbp adjacency of the promoter or in 50kbp of the proximal enhancer or in 500kbp of the distal enhancer. These values were adjustable.

Please note that a DMR might be annotated with enhancers or promoters as well as exons, intronic and intergenic regions of multiple genes. An order of precedence [Promoter/Enhancer/Exons > Intronic > Intergenic] was used to exclude low-priority annotations. For example, if a region is associated with a promoter of a transcript of a gene, its annotation with introns of other transcripts of the same gene was not considered. These annotations were useful, because differential methylation had different effects when it occurred in various regions, and identifying the locations with differential methylation provided crucial insights for disease biomarkers and potential epigenetic therapies.

#### Visualization

To facilitate the analysis of differential methylation, DiffMethylTools offered diverse plots to visualize the methylome, DMLs/DMRs and their annotations as shown in Figure 1:

- **Histogram of coverage**: Coverage histogram was essential for assessing data quality and guiding filtering decisions. Reliable detection of differential methylation required sufficient read depth. Low coverage increased noise and uncertainty, while unusually high coverage might reflect artifacts such as alignment errors or repetitive regions. DiffMethylTools offered coverage histograms to visualize each sample for filtering high and low coverage sites.
- **Volcano plots of DMLs**: Volcano plots of methylation difference versus FDRs provided easy visualization of significant DMLs. Users could use them to check potential issues in DML detection and to quickly identify the most biologically relevant and statistically significant changes between two groups.
- **Manhattan plot of differential methylation**: Manhattan plot provided a genome-wide view of DMLs by plotting statistical significance (–log10 FDR) against chromosome loci. This visualization enabled users to quickly and visually identify chromosomal loci with clustered epigenetic changes. Such plots were widely used in genome-wide association study (GWAS) data or for known disease loci.
- **Annotation pie chart**: Various pie charts were generated for visualizing annotation of DMRs (recommended) and DMLs. DiffMethylTools generated plots for three major annotation categories: the distribution of CpG sites in EPIC array, repeat elements, and functional genomic features (e.g., promoters, enhancers, gene bodies, and intergenic regions). These visualizations helped to identify the overall distribution of differential methylation.
- **Visualization of single DMRs**: DiffMethylTools also provided high-resolution visualizations of individual DMRs, enabling detailed interpretation of locus-specific methylation changes. Two visualization subplots were generated to illustrate per-sample methylation levels or averaged methylation within each group. For the latter subplot, standard deviation within each group was used to illustrate methylation variability. This visualization was also associated with genomic context, overlaying annotated regulatory features (e.g., enhancers) and repetitive elements (e.g., SINEs, LINEs), offering annotation of single DMRs of interest to further investigate biologically meaningful DMRs.
- **Clustering visualization of DMRs in upstream regions of genes**: This clustering view offered an integrative and interpretable summary of how DMRs located in potential promoter regions affected the activities of genes. It enabled users to pinpoint DMR patterns across genes.
- **Gene region vs. methylation difference scatter plots**: DiffMethylTools also generated detailed visualizations that linked differential methylation with specific gene-associated regions. After identifying DMRs, DiffMethylTools grouped methylation levels in 100 bp window, and linked them to intron, exon, upstream, and CCREs using scatter plots. This visualization helped to localize strong methylation shifts in different annotation categories.

**Figure 1:**
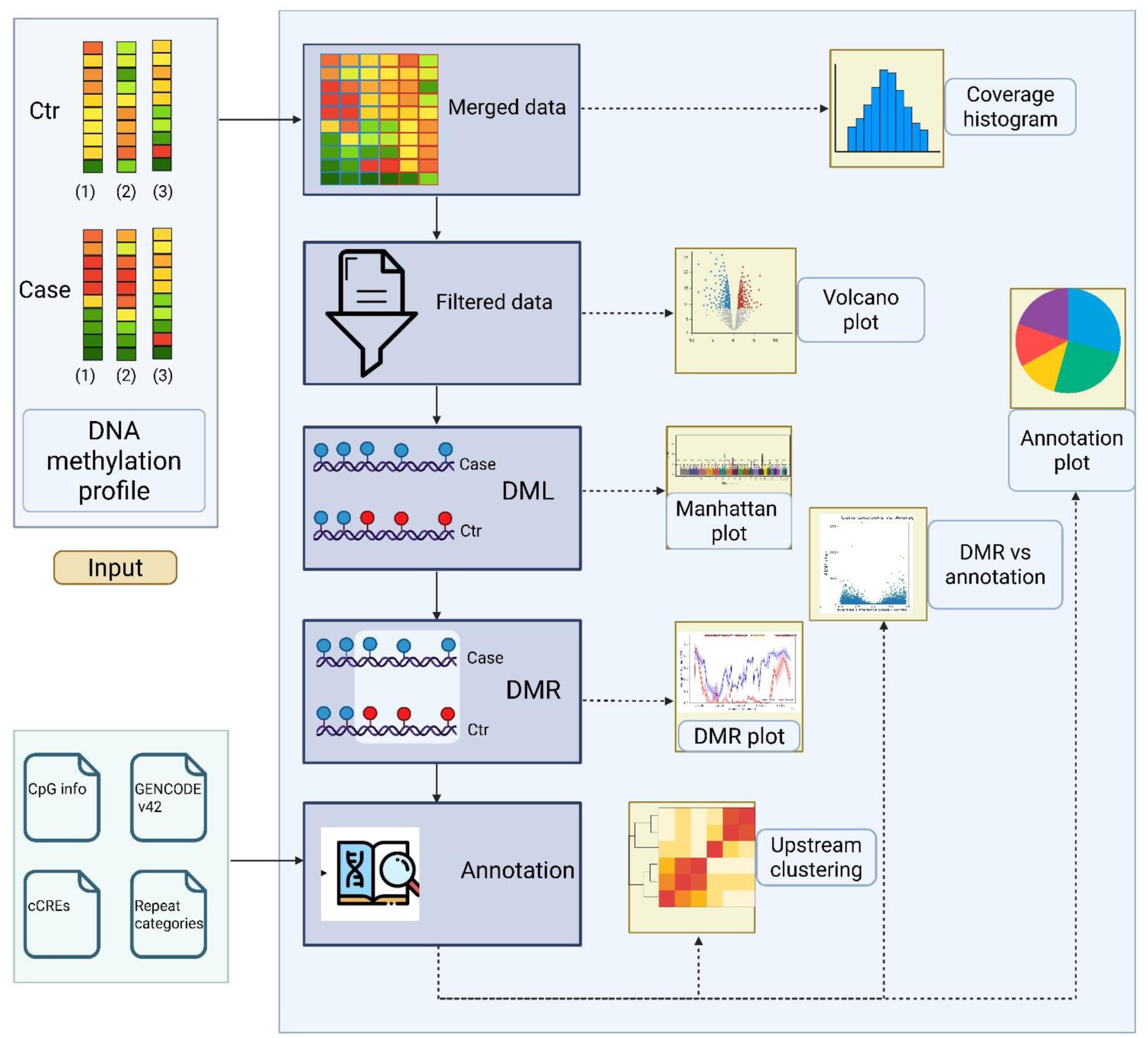
Flowchart of DiffMethylTools. cCRE: candidate cis-Regulatory Elements; DMR: differential methylation regions; DML: differential methylation loci.; (1)-(3): the input methylomes where each row represented CpG site.

### 2.3 State-of-the-art methods for differential methylation detection

To demonstrate the advantage of DiffMethylTools, we evaluated four widely used existing tools on the same set of datasets for differential methylation detection.

*<u>MethylKit</u>* was a comprehensive R package specifically designed to analyze DNA methylation data particularly derived from high-throughput bisulfite sequencing technologies. To detect differential methylation for CpG sites or positions, methylKit used Fisher’s exact test for small sample size datasets or low-coverage datasets, or logistic regression to account for multiple covariates or continuous variables. To detect DMRs, MethylKit employed fixed-size windows to segment genome and then identified whether the windows were differentially methylated. Fixed-size windows were not a good solution to detect DMRs since the size of a DMR varies between tens of bp and thousands of bp. MethylKit offered visualization like histograms of coverage and methylation for individual samples, correlation between samples, and hierarchical clustering and principal component analysis (PCA) to cluster samples. But MethylKit heavily relied on the toolkit genomation^58^ to annotate DMRs with genomic features such as promoters, exons, introns, and intergenic regions and other functional genomic regions.

*<u>DSS (Differential DNA Methylation Analysis for Sequencing)</u>*was another statistical method to identify DMLs and DMRs. It used a Wald test for beta-binomial distributions to compute p-values for DMLs, and then clustered DMLs to DMRs. But DSS does not provide any annotation and visualization functions for detected DMLs and DMRs.

*<u>Bsseq (Bisulfite sequencing analysis)</u>* used t-statistics to detect DMRs. It incorporated the BSmooth algorithm to smooth methylation estimates across CpG sites, reduce technical noise and increase the signal-to-noise ratio. It defined a DMR region by specifying a minimum number of CpG sites in a genomic region. This region-based method is useful for low-coverage data, where raw methylation estimates may be more variable.

*<u>MethylSig</u>* was another specialized R package to detect DMLs and DMRs. It employed a beta-binomial regression framework that accounted for biological replication and overdispersion, enabling more accurate identification of DMLs with modest sample sizes. It detected DMRs by clustering significant DMLs based on genomic proximity, or by fixed windows. MethylSig provided scale-aware visualization for CpG-level data and genomic context in narrow regions, and a simplified overview of features and significance in broader regions.

### 2.4 Evaluation

There were no benchmark DMLs and DMRs in real methylation datasets. We thus provided two ways to evaluate the detection performance of differential methylation. One was based on known DMLs on simulation data. On the simulation data, a position with a Cohen distance of > 0.5 was considered differentially methylated. The Cohen distance was calculated using the equation below:

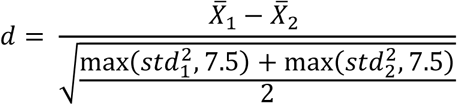

where 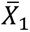 and 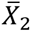 were mean methylation of group 1 and group 2 respectively, and std_1_ and std_2_ were the standard deviation of group 1 and group 2 respectively. A minimum variance threshold (of 7.5) was used to mitigate the impact of unrealistically small variance estimates.

Based on known DMLs, we used recall, precision and F-1 measures to evaluate the detection performance of each tool.

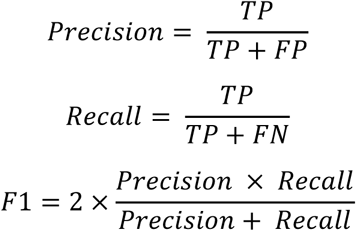

Where TP was the number of known DMLs which were identified as DMLs by a tool, FP was the number of identified DMLs which were not known DMLs, and FN was the number of known DMLs which were not identified as DMLs by a tool.

On real methylation data, we assumed that a DML was more reliable if it was detected by 2+ tools and considered them as benchmark DMLs. We then used the benchmark DMLs to calculate precision, recall and F1 for performance comparison. Please note that we evaluated the performance of five tools, and to avoid bias, a test tool was not used to generate benchmark DMLs.

Similarly, to generate benchmark DMRs for real methylation datasets, we employed a consensus-based strategy. Specifically, when we tested the performance of a tool T1 on a dataset, we identified DMRs that were detected by 3+ of the other four tools as benchmark DMRs. To account for minor discrepancies, nearby benchmark DMRs were merged if they were separated by less than 100 base pairs. The benchmark DMRs were trimmed to ensure that both start and end coordinates corresponded to annotated CpG sites. After that, we compared the benchmark DMRs against detected DMRs to calculate precision, recall and F1 for performance evaluation, where the overlap of benchmark DMRs and detected DMRs was considered as TP, the undetectable regions in benchmark DMRs as FN, and the non-benchmark regions in detected DMRs as FP.

## 3. Results

We evaluated DiffMethylTools and four existing tools using three strategies. We first evaluated their performance on simulation data with known DMLs. We then compared the detection performance of DMLs and DMRs on real data using benchmark DMLs and DMRs. After that, we conducted functional annotation analysis, and discussed visualizations offered by our tool.

### 3.1 Detection performance on simulation data

We run all four tools (bsseq was not included because it cannot detect DMLs) on the simulation data with known DMLs. The volcano plots of detected DMLs were illustrated in Figure 2 (a-d). Different tools had similar range of methylation differences, but their shape of the volcano plots was unique, because they used different statistical models to calculate p-values. DiffMethyTools was the most conservative with fewer DMLs with smaller FDR, while MethylKit and MethylSig were more dispersed. In particular, MethylKit detected a lot of DMLs whose methylation difference was less than 20, but their FDRs were pretty higher---even higher than a lot of DMLs whose methylation difference was much larger. This indicates potential concerns of detecting too many false positives in MethylKit.

**Figure 2:**
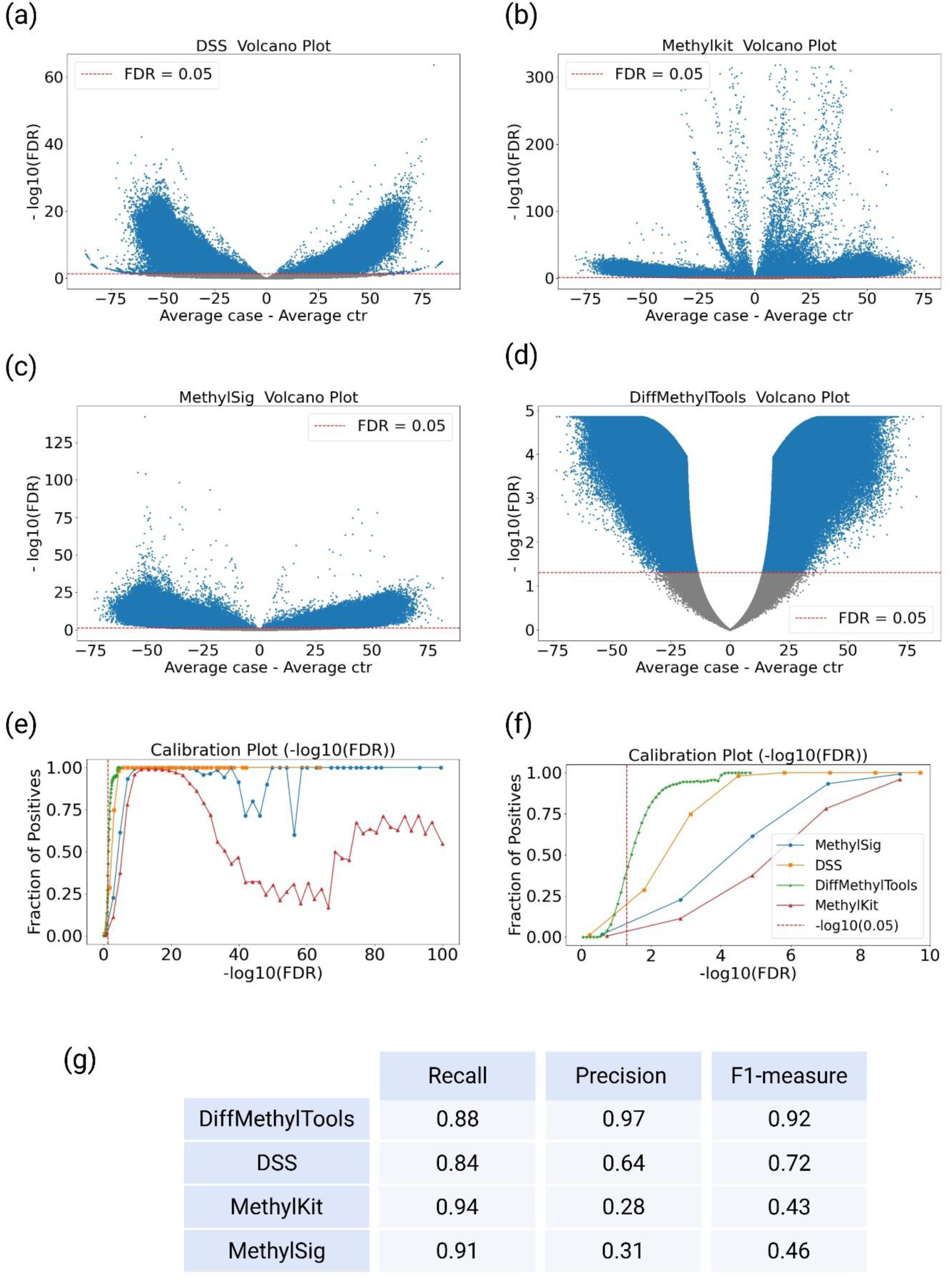
Detection performance of differential methylation loci(DMLs) by four tools on the simulation data. Volcano plots of detected DMLs by methylKit (a), DSS (b), MethylSig (c) and DiffMethylTools (d) where x-axis represented the methylation difference of group mean and y-axis represented the negation of logarithm of corrected p-values (FDR). Each dot denoted a DML, and a horizontal red line indicated the FDR significance threshold of 0.05. The distribution of known DMLs against FDR was shown in (e) for all DMLs and (f) for the DMLs with smaller range of FDR. X-axis represented the negation of logarithm of FDR and y-axis represented the precision of predicted DMLs with FDR smaller than a threshold in x-axis.

To investigate this situation, we used reliability diagrams by plotting the association of the negation of logarithm of FDRs, *i.e.*, -log10(FDR), against observed frequencies of known DML outcomes. In reliability diagrams, a tool with excellent performance should exhibit a low fraction of known DMLs when -log10(FDR) < -log10(0.05) ---reflecting the tool’s ability to avoid false negatives, and large fraction of known DMLs when -log10(FDR) > -log10(0.05)---demonstrating the tool’s effectiveness to accurately identify known DMLs. A perfect model is expected to have a sharp transition around -log10(0.05). In Figure 2 (e) and (f), all tools exhibited an increase in the fraction of known DMLs around -log10(0.05). DiffMethylTools and DSS maintained higher percentage of DMLs as -log10(FDR) become larger. The higher percentage of known DMLs by DiffMethylTools first occurred at approximately x = 4, while that by DSS was around x=5, suggesting that DiffMethylTools was more precisely separate significant and non-significant methylation loci. In contrast, MethylKit and MethylSig reached their maximums with much smaller FDR, and their percentage of DMLs dropped significantly as the FDR become smaller, suggesting that a lot of detected DMLs by MethylKit and MethylSig were not known DMLs even with significantly smaller FDR.

To quantitatively measure the detection performance, we calculated the recall, precision and F1, as shown in Figure 2g. DiffMethylTools achieved higher F1-score (0.92), which was much higher than DSS (0.72), MethylKit (0.43) and MethylSig (0.46). DiffMethylTools’ precision is 0.97, 0.25 higher than DSS, 0.69 higher than methylKit and 0.66 higher than MethylSig, suggesting that the detected DMLs by DiffMethylTools were reproduced by existing tools, while existing tools detected a lot of DMLs which could not be detected by other tools. MethylKit and MethylSig both achieved high recall values (0.94 and 0.91, respectively), but the tradeoff was low precision. Thus, our tool exhibited better performance to detect DMLs. Overall, DiffMethylTools demonstrated higher capabilities to filter out false DMLs while maintaining higher recall rates.

### 3.2 Evaluation of DML detection on real methylation data

We then evaluated the detection performance of DMLs on three long-read methylation data and two WGBS data. The F1 scores were summarized in Table 1 for the four tools DiffMethylTools, DSS, MethylKit and MethylSig, and the detail was described in Supplementary Tables 1-5.

**Table 1:** F1 scores of the detection of differential methylation loci (DMLs) across five datasets. ONT: Oxford Nanopore sequencing; WGBS: whole-genome bisulfite sequencing.

|  |  | DiffMethylTools | DSS | MethylKit | MethylSig |
| --- | --- | --- | --- | --- | --- |
| ONT data | HCT116 | 0.91 | 0.85 | 0.66 | 0.79 |
|  | MO/ND | 0.48 | 0.56 | 0.29 | 0.33 |
|  | AD | 0.15 | 0.05 | < 0.01 | 0.04 |
| WGBS | Liver cancer | 0.44 | 0.55 | < 0.01 | 0.28 |
|  | NK/B cells | 0.78 | 0.88 | 0.42 | 0.6 |

Across all five datasets, DiffMethylTools consistently demonstrated superior performance compared to MethylKit and MethylSig. On the HCT116 dataset, DiffMethylTools achieved the highest F1 score (0.91), significantly surpassing MethylKit (0.66) and MethylSig (0.79). On the two cell-type datasets, MO/ND and NK/B cells, DiffMethylTools outperformed MethylKit and MethylSig by 0.15-0.19 on MO/ND, and by 0.18-0.36 on NK/B cells. On the two methylation datasets with real human diseases (Alzheimer’s disease and liver cancer), the F1-score of MethylKits and MethylSig was much worse than those achieved by DiffMethylTools. On the Liver cancer dataset, DiffMethylTools acheived an F1 score of 0.46, markedly higher than MethylKit (< 0.01) and MethylSig (0.28). On the AD dataset, DiffMethylTools achieved a modest F1 score (0.15), outperforming DSS (0.05), MethylSig (0.04), and MethylKit (< 0.01). The poor performance of MethylSig was because a lot of detected DMLs by MethylSig could not be reproduced by other tools (Supplementary Tables 1-5). Compared to DSS, DiffMethylTools achieved overall better DML detection on long-read sequencing data and comparable performance across the five datasets. However, as we discussed below, DSS generated much worse performance to detect DMRs than DiffMethylTools.

#### Qualitative evaluation of DMLs

To investigate the potential issues of DML detection, we visualized DMLs by each tool on the HCT116 data using volcano plots in Figure 3 with methylation difference (Δβ) for x-axis and –log₁₀(FDR) as y-axis.

**Figure 3:**
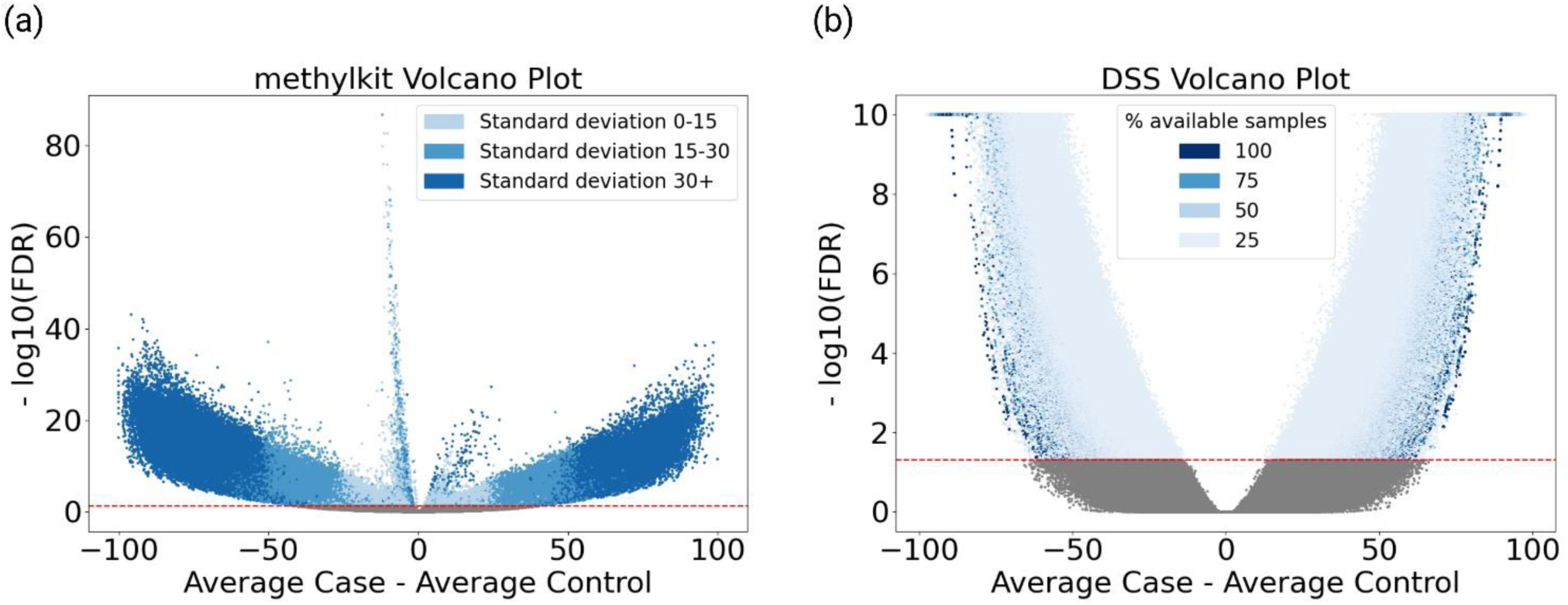
Volcano plots of DMLs detected by methylKit (a) and DSS (b) on HCT116 data. On (a), DMLs were colored based on their standard deviations, with dark blue indicating high standard deviations and lighter shades representing lower variability. On (b), DMLs were colored according to how many of the four available samples had enough coverage support for DMLs, with all samples supported (i.e., all four) in dark blue, and less samples in lighter shades.

As shown in Figure 3a and b, MethylKit and MethylSig detected a lot of DMLs. MethylKit detected a lot of DMLs with smaller Δβ or smaller standard deviation of methylation levels across all samples in the dataset. It is usually hard to distinguish a DMLs with smaller Δβ or smaller standard deviation from random changes, and thus those DMLs contained more noises. On the other hand, DSS detected a lot of DMLs whose coverage was lower in samples (Figure 3b).

To further investigate the difference between detected DMLs via our tool and existing tools, we plotted the methylation difference and standard deviation of different sets of detected DMLs by different tools in Figure 4. The different sets of DMLs included three sets of DMLs identified by each of the three state-of-the-art tools (SOTA) but not by DiffMethylTools (e.g., DSS − DiffMethylTools), (2) three sets of DMLs shared between DiffMethylTools and each of the three SOTA tools (e.g., DiffMethylTools ∩ DSS), (3) a set of DMLs identified by all SOTA methods (intersection) but not by DiffMethylTools (SOTA − DiffMethylTools), and (4) a set of DMLs uniquely detected by DiffMethylTools (DiffMethylTools − SOTA).

**Figure 4:**
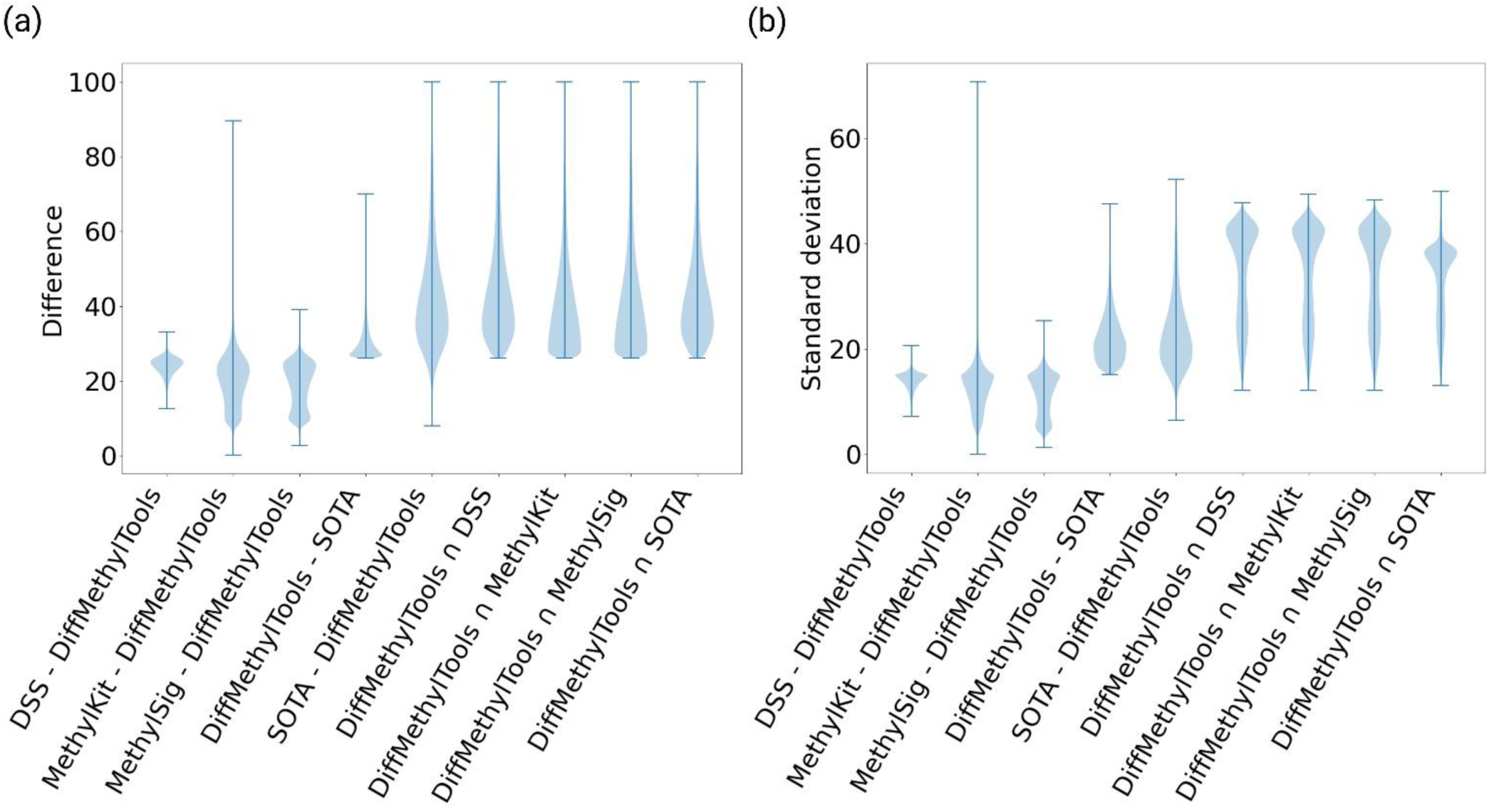
Violin plots of the absolute difference (a) and the standard deviation (b) of DMLs identified by different methods. The format of X-axis “T1 − T2” was for DMLs detected by T1 but not T2, or “T1 ∩ T2” for DMLs detected by both T1 and T2. SOTA: the intersection of DMLs detected by three state-of-the-art (SOTA) methods.

As shown in Figure 4, the DMLs identified solely by DiffMethylTools, the DMLs detected by the intersection of the three SOTA tools, and the overlap DMLs of DiffMethylTools and existing tools had similar level of larger methylation difference (Figure 4a) than the DMLs detected only by each of the three SOTA tools. This suggested that DiffMethylTools’ detection was more reliable. Also, the overlap DMLs of DiffMethylTools and existing tools had larger standard deviation than the DMLs detected by the intersection of the three SOTA tools and the DMLs identified solely by DiffMethylTools. In contrast, the DMLs detected only by each of the three SOTA tools had the smallest standard deviations. Since the standard deviation was calculated for all samples across groups, smallest standard deviations suggested less difference between groups. Thus, these pieces of evidence suggested that the detected DMLs by DiffMethylTools was more reliable.

In summary, different tools used various statistical models and detected DMLs with various characteristics. But our proposed method DiffMethylTools exhibited better capability to capture greater methylation differences and higher standard deviations in DMLs, and thus achieved overall better performance.

### 3.3 Investigate DMR detection on real methylation data

#### 3.3.1 DMR detection performance

We also evaluated DMR detection for five tools, i.e., DiffMethylTools, DSS, MethylKit, MethylSig and bsseq, on the five datasets. To benchmark these tools, we found the overlap DMR regions detected by 3+ tools on a dataset as the benchmark DMRs (as described in the Methods section). These benchmark DMRs on each data were used to calculate precision, recall and F1-score of each tool. In real-world applications, a subset of differential methylation patterns would usually be selected as downstream inputs, and thus, it was very important to get a higher precision set of DMRs. Thus, we investigated the precision of each method in Table 2, and reported recall and F1 scores in (Supplementary Tables 6-10).

**Table 2.** Precision of DMR detection by the five tools on the three ONT data and two WGBS data. ONT: Oxford Nanopore sequencing; WGBS: whole-genome bisulfite sequencing.

|  |  | DiffMethylTools | DSS | MethylKit | MethylSig | bsseq |
| --- | --- | --- | --- | --- | --- | --- |
| ONT data | HCT116 | **0.68** | 0.61 | 0.09 | 0.06 | 0.64 |
|  | MO/ND | **0.84** | < 0.01 | < 0.01 | < 0.01 | 0.1 |
|  | AD | **0.21** | 0.07 | < 0.01 | < 0.01 | 0.01 |
| WGBS | Liver cancer | 0.13 | < 0.01 | < 0.01 | 0.02 | **0.19** |
|  | NK/B cells | 0.71 | 0.74 | 0.02 | 0.04 | **0.84** |

As shown in Table 2, DiffMethylTools consistently outperformed the other methods on long-read data. For example, on HCT116, the precision of DiffMethylTools was 0.68, 0.07 higher than DSS, 0.59 higher than methylKit, 0.62 higher than MethylSig and 0.04 higher than bsseq. On the AD dataset, DiffMethylTools’ precision (0.21) was 0.14 higher than DSS and 0.2 higher than methylKit, MethylSig and bsseq. Similarly, on the MO/ND data, DiffMethylTools’ precision was 0.84, which was 0.74 higher than bsseq and 0.83 higher than DSS, methylKit, and MethylSig. On the two short-read methylation data, DiffMethylTools performance was lower than the best-performed existing tool bsseq, but still ranked as the second or third best positions. These results demonstrated that DiffMethylTools achieved overall better detection performance for DMRs and highlighted the robustness and generalizability of DiffMethylTools across different sequencing platforms and biological conditions.

### 3.4 Functional annotation analysis of DMRs

#### 3.4.1 The distribution of functional annotations

DiffMethylTools offered a comprehensive annotation for DMLs and DMRs by intersecting methylation patterns with various genomic annotations as well as interpretable visual summaries for genomic feature distributions with two types of visualization: a site-based visualization and an annotation-based visualization. A site-based visualization counted the number of CpG sites in methylation patterns that overlapped with an annotation category and thereby reflected the relative concentration or density of methylation signal within different genomic features. In contrast, an annotation-based visualization counted the occurrence of genome annotation that overlapped with methylation patterns. This dual-level resolution revealed notable differences in interpretation.

We illustrated these annotations using the DMRs detected on NK/B cells in Figure *5* (for site-based visualization) and in Figure 6 (annotation-based visualization). We found that 92.3% of sites in DMRs were not detectable in EPIC array (Figure *5*a), whereas 54.2% of detected DMRs were completely missed when using EPIC array (Figure 6a). Both suggested that a lot of differential methylation would be missed if EPIC array data were used. Figure *5*b demonstrated that ∼40% CpG sites in DMRs and ∼70% of DMRs were located in repeat regions, and these patterns could not be reliably detected if short-read sequencing data were used. In Figure *5*c and Figure *5*d as well as Figure 6c and Figure 6d, ∼53-56% of DMRs were located in enhancers and ∼15-18% in exons, while ∼5-6% in intergenic regions. Enhancers and exons were enriched in DMRs compared to that ∼7% of hg38 were defined as cCRE elements while 2% for exons. In contrast, intergenic regions (65% of hg38) were depleted. These annotations indicated that the detected DMRs were biologically meaningful. Our tool DiffMethylTools facilitated biological interpretation.

**Figure 5.**
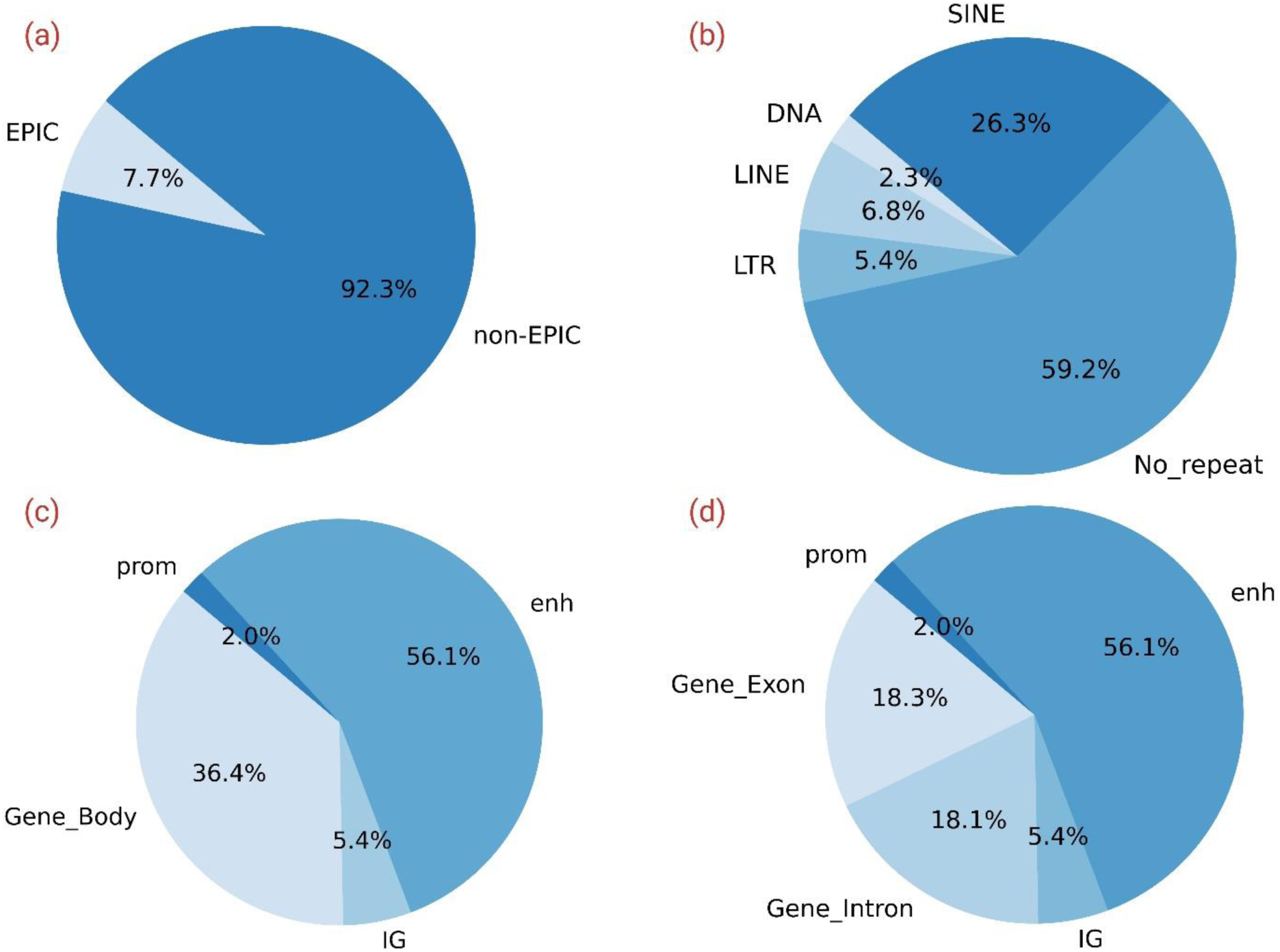
Site-based visualization of DMRs and genomic annotations on NK and B cells. Pie charts display the proportional CpG distribution overlapped DMRs in various genomic regions: CpG sites in EPIC vs. non-EPIC array (a), repeat elements (b), and functional annotations (promoters, enhancers, gene body, intergenic) in (c) and (d).

**Figure 6.**
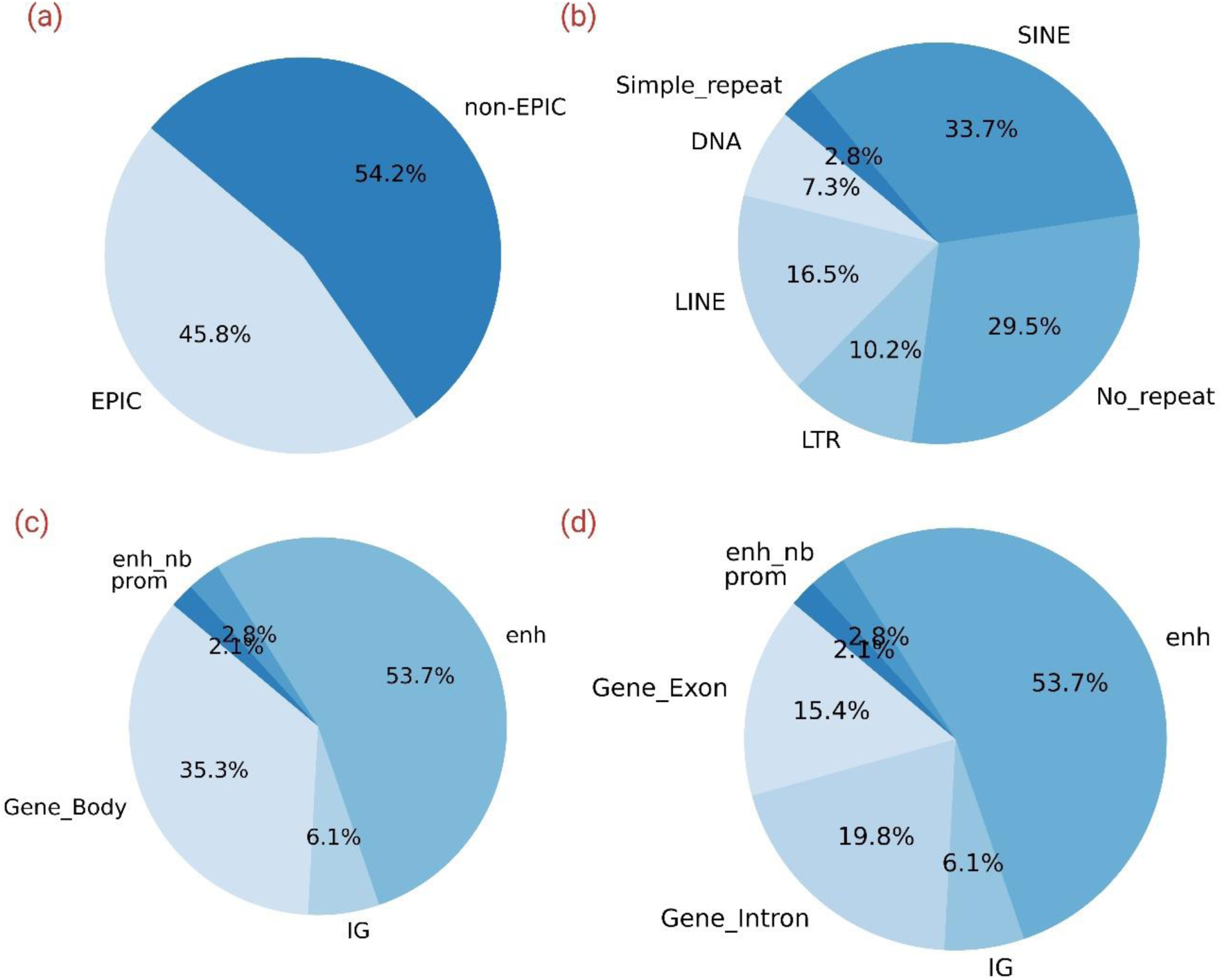
Annotation-based visualization of DMRs between NK and B cells. CpG sites in EPIC vs. non-EPIC array (a), repeat elements (b), and functional annotations (promoters, enhancers, gene body, intergenic) in (c) and (d).

#### 3.4.2 Annotation of DMRs

Besides the visualization of annotation distribution, DiffMethylTools generated high-resolution visualization of each DMR. An example of such visualization was illustrated in Figure *7* for a DMR (chr1:3664875–3667889) detected on the NK/B cells. This visualization not only contained per-sample methylation levels in the DMR (Figure *7*a) but also included group-level summary (Figure *7*b). Figure *7*a revealed a consistent hypermethylation pattern in NK cells compared to B cells, while Figure *7*b supported this observation with smaller intra-group variability in both conditions, indicating strong reproducibility across replicates. In addition, genomic features and repeat categories that overlapped with the DMR were also illustrated, enabling users to prioritize biologically meaningful DMRs for follow-up analysis and supporting hypothesis-driven exploration of epigenetic regulation. For example, this DMR overlapped an annotated enhancer element (ENSR1_BPMG^59^ overlapped with EH38E1312908, EH38E1312909 and EH38E1312910 in Figure *7*), which was known to be active in B cells but inactive in NK cells, corroborating the observed methylation differences. These features suggested that this DMR may play a biologically meaningful role in cell-type-specific gene regulation. This integrative visualization clearly demonstrated how DiffMethylTools contextualized epigenetic variation within the broader regulatory and structural genome, facilitating downstream investigation.

**Figure 7.**
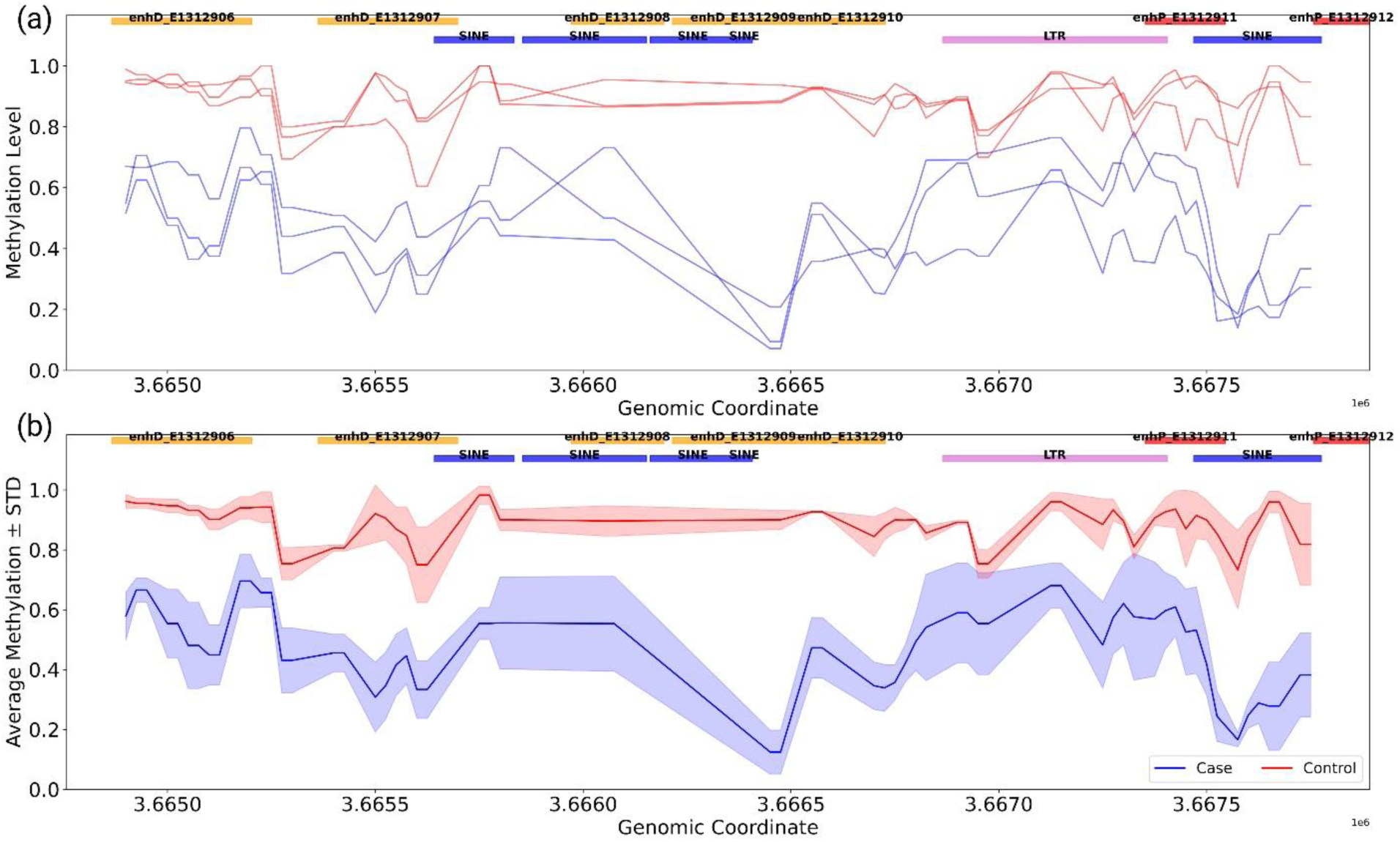
Visualization of an example differentially methylated region (DMR) on chr1:3664875-3667889 identified on NK/B dataset (with NK cells (control) and B cells (case)) by DiffMethylTools. (a): methylation levels for each sample separately; (b): the methylation mean in each group with standard deviation. SINE: Short interspersed nuclear elements; LTR: Long Terminal Repeat; EnhP: proximal enhancer with its identifier; enhD: distal enhancer with its identifier.

### 3.4 Visualization

DiffMethylTools offered diverse visualizations to simplify methylation investigation, as discussed in the Methods section. We discussed several visualizations below and more visualization such as per-sample coverage plots could be found in Supplementary Figures 1-5.

One important visualization was to cluster upstream regions of genes which contained DMRs. This clustering view offered an integrative and interpretable summary of regulatory methylation changes proximally to gene starts and was useful to conduct cross-gene investigation of DMRs. An example was illustrated Figure 8 for the dataset of NK/B cells. As shown in Figure 8, genes’ upstream with hypomethylation and hypermethylation was clustered separately. Also, hypomethylation upstream regions were further clustered according to the location of the hypomethylation, suggesting that some genes tend to be hypomethylated in similar relative locations of their upstream regions. This visualization aided in detecting coordinated methylation patterns across gene-proximal regions, exploring how epigenetic shifts might drive transcriptional regulation.

**Figure 8.**
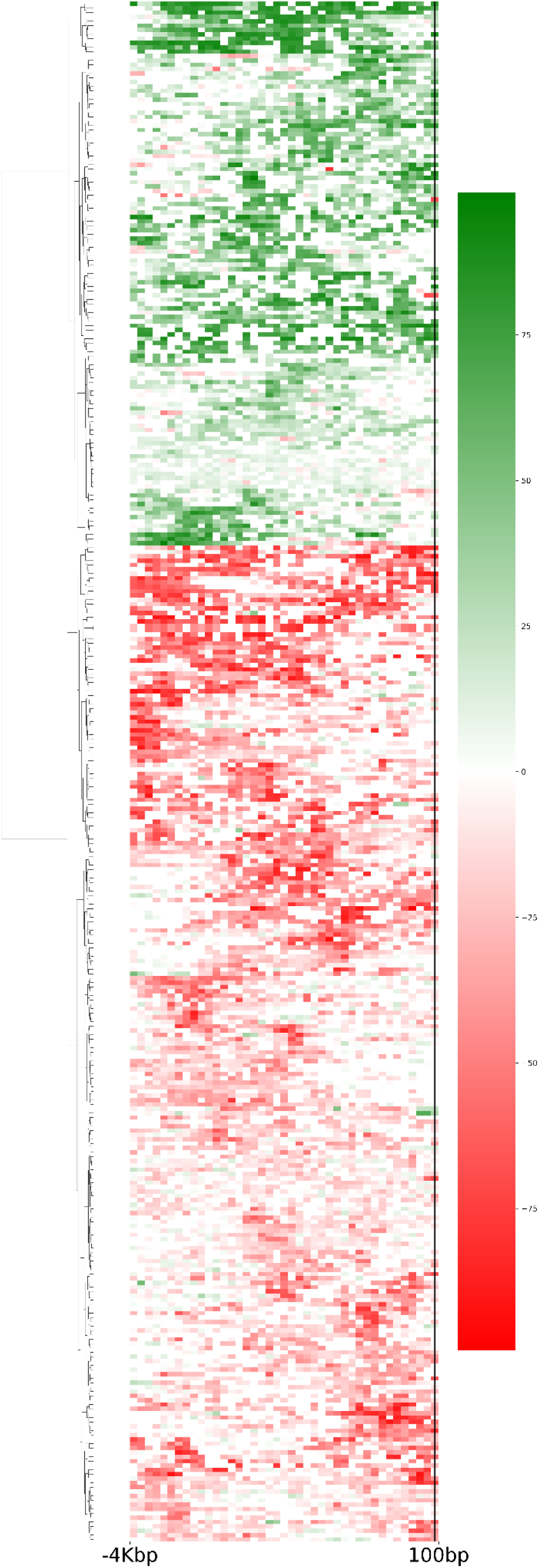
Clustering of upstream regions of genes that overlapped differentially methylated regions (DMRs) in NK/B cell dataset. Each row represented a gene whose upstream contained 1+ DMRs, and each column corresponded to a 100bp window, and color intensity indicated the methylation difference between two groups after averaging methylation levels for a 100bp window: red for hypomethylation and green for hypermethylation.

Besides the clustering of upstream, DiffMethylTools also generated scatterplots of differential methylation patterns against introns, exons, upstream regions, and CCRE regions. An example was shown in Figure 9 for DMRs detected on the dataset of NK/B cells. It seemed that introns tended to exhibit more hypermethylation or hypomethylation sites with larger methylation differences, and CCRE regions tended to contain less CpG sites. The methylation change behaviors were similar for exon and upstream regions: diverse ranges (from moderate changes to larger deviation) of methylation changes but with less CpG sites than intron regions. These results demonstrated that methylation changes in various genomic regions were different. DiffMethylTools outputted this information in a csv file that users could find genes of interest.

**Figure 9:**
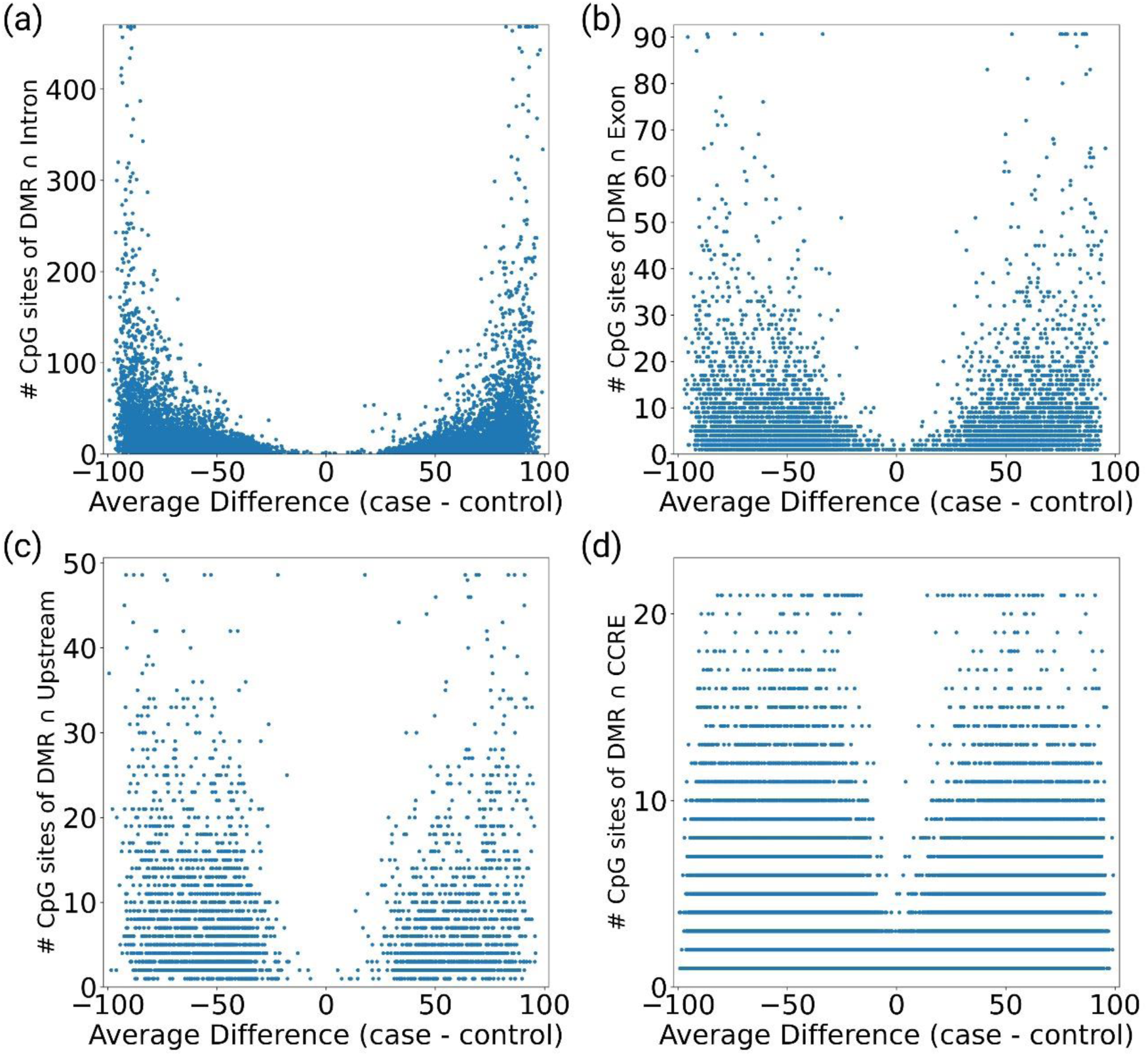
Scatter plots of genomic regions against the number of CpG sites in DMRs detected on the dataset of NK/B cells for four genomic features: introns (a), exons (b), upstream regions (c), and candidate cis-regulatory elements (CCREs) in (d). x-axis: the average methylation difference; y-axis: the number of CpG sites in DMRs per gene region (y-axis).

Another helpful visualization was the Manhattan plot for the detected DMLs across the genome and one example was presented in Figure *10* for the MO/ND dataset. This plot helped users to quickly visualize the detected DMLs and potential DMLs clusters. Please note that this plot might not be used if there were too many detected DMLs.

**Figure 10:**
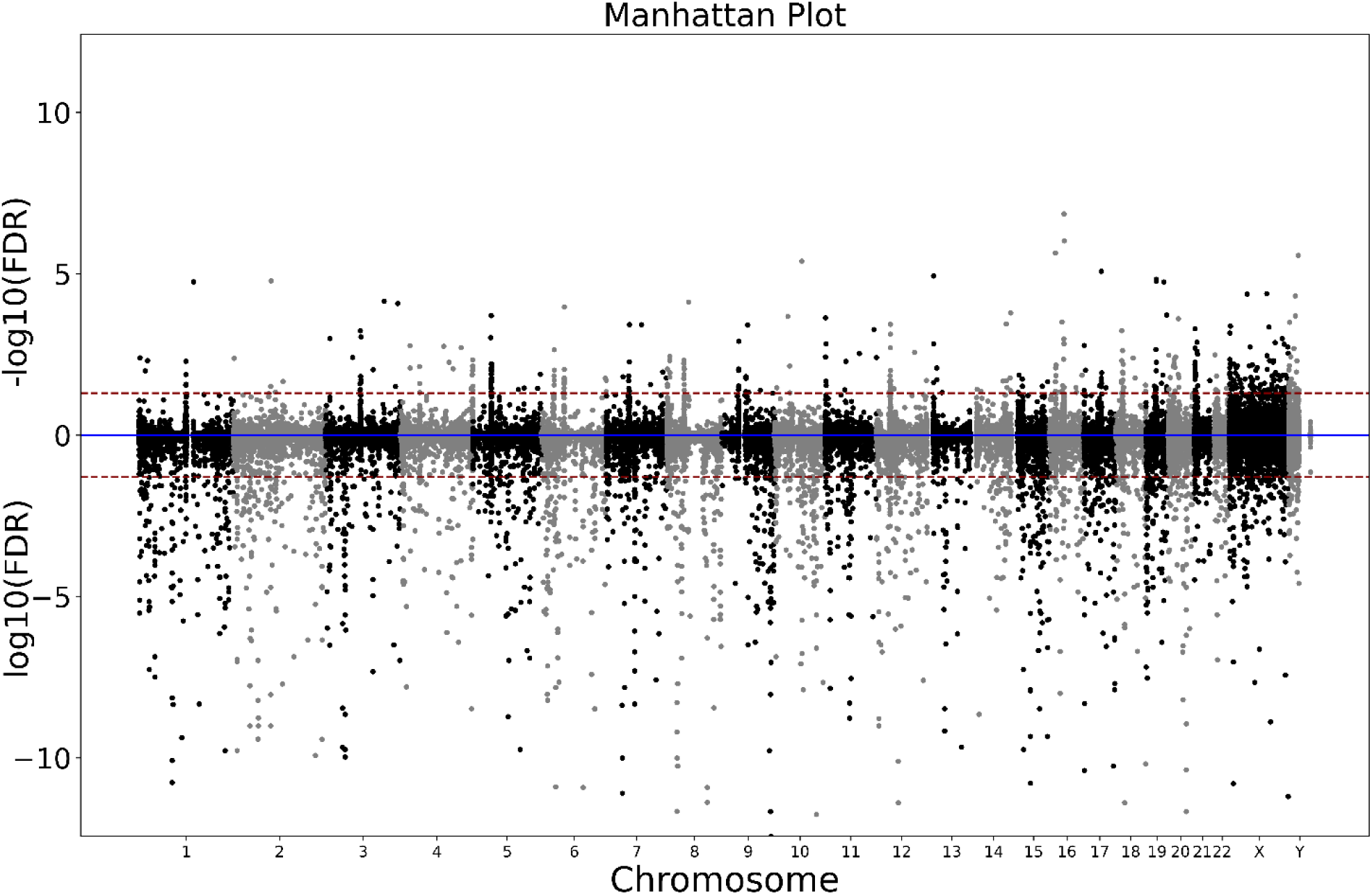
Manhattan plot of DMLs detected on the dataset of the MO/ND dataset. Y-axis: the negative logarithm of FDR (top panel) and the logarithm of FDR (bottom panel); x-axis: the chromosome and genomic coordinates.

## 4. Discussion

Differential methylation biomarkers were critical to investigating epigenetic contributions in various downstream analysis. We introduced DiffMethylTools, a comprehensive framework to facilitate this investigation by integrating the detection, annotation, and visualization of differentially methylated loci (DMLs) and regions (DMRs) in a single command. DiffMethylTools enhanced robust detection of differential methylation compared to existing detection methods and achieved better performance on long read methylomes. Volcano plots and calibration curves highlighted DiffMethylTools’ ability to clearly separate signal from noise, resulting in high-confidence DML detection with minimal false positives.

Another important feature of DiffMethylTools was its ability to annotate methylation changes with comprehensive genomic context as well as various visualizations. DiffMethylTools mapped DMLs and DMRs to regulatory features, gene elements, and repetitive regions, enabling deeper biological interpretation. This capability was particularly useful to link methylation differences with potential functional effects. Visualizations such as Manhattan plots, gene-centric scatterplots, and per-region methylation heatmaps made DiffMethylTools accessible to both computational and experimental biologists to illustrate and interpret differential methylation.

Nonetheless, DiffMethylTools could be improved in future. First, DiffMethylTools detected DMLs before clustering them to DMRs. This single site detection may be sensitive to outliers in small sample sizes. It is better to integrate local co-methylation observation^52–55^ to improve the robustness of the DML detection. Second, the annotation analysis was based on hg38. Annotation for telomere-to-telomere reference genomes needs to be integrated when corresponding resources become available.

## 5. Conclusion

In summary, DiffMethylTools is a robust, interpretable, and user-friendly solution for differential methylation investigation across sequencing technologies. It streamlined epigenomic research by robust detection, optimized support for long-read data, advanced annotation features, and integrated visualization capabilities. Its excellent detection performance across both synthetic and real data highlighted its potential as a valuable tool for epigenomic research and clinical methylation studies.

## Funding

This research was funded by the National Institute of General Medical Sciences grant number P20GM121325 and a UNLV startup grant. The funders had no role in the study design, data collection and analysis, the decision to publish, or the preparation of the manuscript.

## Author Contributions

Q.L. conceived and supervised the study, Q. L., H.D. and E. K. implemented the pipeline. H.D. and E. K. conducted the data analysis and tested the pipeline. J. W. provided the ND/MO cell samples. H. D. and Q.L. wrote the manuscript. All authors have read, revised and agreed to the published version of the manuscript.

## Competing interests

The authors declare no competing interests.

## Data and code availability

The codes are available on GitHub via https://github.com/qgenlab/DiffMethylTools.

## Notes

### Competing Interest Statement

The authors have declared no competing interest.

## References

1. Bird, A. (2002). DNA methylation patterns and epigenetic memory. Genes Dev. 16, 6–21. 10.1101/gad.947102.

2. Chowdhury, B., Cho, I.-H., Hahn, N., and Irudayaraj, J. (2014). Quantification of 5-methylcytosine, 5-hydroxymethylcytosine and 5-carboxylcytosine from the blood of cancer patients by an enzyme-based immunoassay. Anal. Chim. Acta 852, 212–217. 10.1016/j.aca.2014.09.020.

3. Elhamamsy, A.R. (2017). Role of DNA methylation in imprinting disorders: an updated review. J. Assist. Reprod. Genet. 34, 549–562. 10.1007/s10815-017-0895-5.

4. Sharp, A.J., Stathaki, E., Migliavacca, E., Brahmachary, M., Montgomery, S.B., Dupre, Y., and Antonarakis, S.E. (2011). DNA methylation profiles of human active and inactive X chromosomes. Genome Res. 21, 1592–1600. 10.1101/gr.112680.110.

5. Hollister, J.D., and Gaut, B.S. (2009). Epigenetic silencing of transposable elements: A trade-off between reduced transposition and deleterious effects on neighboring gene expression. Genome Res. 19, 1419–1428. 10.1101/gr.091678.109.

6. Besselink, N., Keijer, J., Vermeulen, C., Boymans, S., de Ridder, J., van Hoeck, A., Cuppen, E., and Kuijk, E. (2023). The genome-wide mutational consequences of DNA hypomethylation. Sci. Rep. 13, 6874. 10.1038/s41598-023-33932-3.

7. Booth, M.J., Ost, T.W.B., Beraldi, D., Bell, N.M., Branco, M.R., Reik, W., and Balasubramanian, S. (2013). Oxidative bisulfite sequencing of 5-methylcytosine and 5-hydroxymethylcytosine. Nat. Protoc. 8, 1841–1851. 10.1038/nprot.2013.115.

8. Porreca, G.J. (2010). Genome sequencing on nanoballs. Nat. Biotechnol. 28, 43–44. 10.1038/nbt0110-43.

9. Thompson, J.F., and Steinmann, K.E. (2010). Single Molecule Sequencing with a HeliScope Genetic Analysis System. Curr. Protoc. Mol. Biol. 92. 10.1002/0471142727.mb0710s92.

10. Eid, J., Fehr, A., Gray, J., Luong, K., Lyle, J., Otto, G., Peluso, P., Rank, D., Baybayan, P., Bettman, B., et al. (2009). Real-Time DNA Sequencing from Single Polymerase Molecules. 323.

11. Sereika, M., Kirkegaard, R.H., Karst, S.M., Michaelsen, T.Y., Sørensen, E.A., Wollenberg, R.D., and Albertsen, M. (2022). Oxford Nanopore R10.4 long-read sequencing enables the generation of near-finished bacterial genomes from pure cultures and metagenomes without short-read or reference polishing. Nat. Methods 19, 823–826. 10.1038/s41592-022-01539-7.

12. Lin, B., Hui, J., and Mao, H. (2021). Nanopore Technology and Its Applications in Gene Sequencing. Biosensors 11, 214. 10.3390/bios11070214.

13. Mazzone, R., Zwergel, C., Artico, M., Taurone, S., Ralli, M., Greco, A., and Mai, A. (2019). The emerging role of epigenetics in human autoimmune disorders. Clin. Epigenetics 11, 34. 10.1186/s13148-019-0632-2.

14. Lakshminarasimhan, R., and Liang, G. (2016). The Role of DNA Methylation in Cancer. In DNA Methyltransferases - Role and Function Advances in Experimental Medicine and Biology., A. Jeltsch and R. Z. Jurkowska, eds. (Springer International Publishing), pp. 151–172. 10.1007/978-3-319-43624-1_7.

15. Shireby, G., Dempster, E.L., Policicchio, S., Smith, R.G., Pishva, E., Chioza, B., Davies, J.P., Burrage, J., Lunnon, K., Seiler Vellame, D., et al. (2022). DNA methylation signatures of Alzheimer’s disease neuropathology in the cortex are primarily driven by variation in non-neuronal cell-types. Nat. Commun. 13, 5620. 10.1038/s41467-022-33394-7.

16. Law, P.-P., and Holland, M.L. (2019). DNA methylation at the crossroads of gene and environment interactions. Essays Biochem. 63, 717–726. 10.1042/EBC20190031.

17. Gao, F., Liu, X., Wu, X.-P., Wang, X.-L., Gong, D., Lu, H., Xia, Y., Song, Y., Wang, J., Du, J., et al. (2012). Differential DNA methylation in discrete developmental stages of the parasitic nematode Trichinella spiralis. Genome Biol. 13, R100. 10.1186/gb-2012-13-10-r100.

18. Lv, J., Liu, H., Su, J., Wu, X., Liu, H., Li, B., Xiao, X., Wang, F., Wu, Q., and Zhang, Y. (2012). DiseaseMeth: a human disease methylation database. Nucleic Acids Res. 40, D1030–D1035. 10.1093/nar/gkr1169.

19. Hu, S., Tao, J., Peng, M., Ye, Z., Chen, Z., Chen, H., Yu, H., Wang, B., Fan, J.-B., and Ni, B. (2023). Accurate detection of early-stage lung cancer using a panel of circulating cell-free DNA methylation biomarkers. Biomark. Res. 11, 45. 10.1186/s40364-023-00486-5.

20. Zhang, L., Li, D., Du, F., Huang, H., Yuan, C., Fu, J., Sun, S., Tian, T., Liu, X., Sun, H., et al. (2021). A panel of differentially methylated regions enable prognosis prediction for colorectal cancer. Genomics 113, 3285–3293. 10.1016/j.ygeno.2021.07.010.

21. Yang, T., Li, C., Wei, Q., Pang, D., Cheng, Y., Huang, J., Lin, J., Xiao, Y., Jiang, Q., Wang, S., et al. (2024). Genome-wide DNA methylation analysis related to ALS patient progression and survival. J. Neurol. 271, 2672–2683. 10.1007/s00415-024-12222-6.

22. Tang, J., Xiong, Y., Zhou, H.-H., and Chen, X.-P. (2014). DNA methylation and personalized medicine. J. Clin. Pharm. Ther. 39, 621–627. 10.1111/jcpt.12206.

23. Plant, D., Webster, A., Nair, N., Oliver, J., Smith, S.L., Eyre, S., Hyrich, K.L., Wilson, A.G., Morgan, A.W., Isaacs, J.D., et al. (2016). Differential Methylation as a Biomarker of Response to Etanercept in Patients With Rheumatoid Arthritis. Arthritis Rheumatol. 68, 1353–1360. 10.1002/art.39590.

24. Gaspar, J.M., and Hart, R.P. (2017). DMRfinder: efficiently identifying differentially methylated regions from MethylC-seq data. BMC Bioinformatics 18, 528. 10.1186/s12859-017-1909-0.

25. Li, S., Garrett-Bakelman, F.E., Akalin, A., Zumbo, P., Levine, R., To, B.L., Lewis, I.D., Brown, A.L., D’Andrea, R.J., Melnick, A., et al. (2013). An optimized algorithm for detecting and annotating regional differential methylation. BMC Bioinformatics 14, S10. 10.1186/1471-2105-14-S5-S10.

26. Akalin, A., Kormaksson, M., Li, S., Garrett-Bakelman, F.E., Figueroa, M.E., Melnick, A., and Mason, C.E. (2012). methylKit: a comprehensive R package for the analysis of genome-wide DNA methylation profiles. Genome Biol. 13, R87. 10.1186/gb-2012-13-10-r87.

27. Park, Y., Figueroa, M.E., Rozek, L.S., and Sartor, M.A. (2014). MethylSig: a whole genome DNA methylation analysis pipeline. Bioinformatics 30, 2414–2422. 10.1093/bioinformatics/btu339.

28. Wu, H., Xu, T., Feng, H., Chen, L., Li, B., Yao, B., Qin, Z., Jin, P., and Conneely, K.N. (2015). Detection of differentially methylated regions from whole-genome bisulfite sequencing data without replicates. Nucleic Acids Res., gkv715. 10.1093/nar/gkv715.

29. Feng, H., Conneely, K.N., and Wu, H. (2014). A Bayesian hierarchical model to detect differentially methylated loci from single nucleotide resolution sequencing data. Nucleic Acids Res. 42, e69–e69. 10.1093/nar/gku154.

30. Park, Y., and Wu, H. (2016). Differential methylation analysis for BS-seq data under general experimental design. Bioinformatics 32, 1446–1453. 10.1093/bioinformatics/btw026.

31. Catoni, M., Tsang, J.M., Greco, A.P., and Zabet, N.R. (2018). DMRcaller: a versatile R/Bioconductor package for detection and visualization of differentially methylated regions in CpG and non-CpG contexts. Nucleic Acids Res. 10.1093/nar/gky602.

32. Sun, D., Xi, Y., Rodriguez, B., Park, H.J., Tong, P., Meong, M., Goodell, M.A., and Li, W. (2014). MOABS: model based analysis of bisulfite sequencing data.

33. Dolzhenko, E., and Smith, A.D. (2014). Using beta-binomial regression for high-precision differential methylation analysis in multifactor whole-genome bisulfite sequencing experiments. BMC Bioinformatics 15, 215. 10.1186/1471-2105-15-215.

34. Stockwell, P.A., Chatterjee, A., Rodger, E.J., and Morison, I.M. (2014). DMAP: differential methylation analysis package for RRBS and WGBS data. Bioinformatics 30, 1814–1822. 10.1093/bioinformatics/btu126.

35. Assenov, Y., Müller, F., Lutsik, P., Walter, J., Lengauer, T., and Bock, C. (2014). Comprehensive analysis of DNA methylation data with RnBeads. Nat. Methods 11, 1138–1140. 10.1038/nmeth.3115.

36. Müller, F., Scherer, M., Assenov, Y., Lutsik, P., Walter, J., Lengauer, T., and Bock, C. (2019). RnBeads 2.0: comprehensive analysis of DNA methylation data. Genome Biol. 20, 55. 10.1186/s13059-019-1664-9.

37. Warden, C.D., Lee, H., Tompkins, J.D., Li, X., Wang, C., Riggs, A.D., Yu, H., Jove, R., and Yuan, Y.-C. (2013). COHCAP: an integrative genomic pipeline for single-nucleotide resolution DNA methylation analysis. Nucleic Acids Res. 41, e117–e117. 10.1093/nar/gkt242.

38. Hansen, K.D., Langmead, B., and Irizarry, R.A. (2012). BSmooth: from whole genome bisulfite sequencing reads to differentially methylated regions. Genome Biol. 13, R83. 10.1186/gb-2012-13-10-r83.

39. Hebestreit, K., Dugas, M., and Klein, H.-U. (2013). Detection of significantly differentially methylated regions in targeted bisulfite sequencing data. Bioinformatics 29, 1647–1653. 10.1093/bioinformatics/btt263.

40. Piao, Y., Xu, W., Park, K.H., Ryu, K.H., and Xiang, R. (2021). Comprehensive Evaluation of Differential Methylation Analysis Methods for Bisulfite Sequencing Data. Int. J. Environ. Res. Public. Health 18, 7975. 10.3390/ijerph18157975.

41. Rabiner, L., and Juang, B. (1986). An introduction to hidden Markov models. IEEE ASSP Mag. 3, 4–16. 10.1109/MASSP.1986.1165342.

42. Yu, X., and Sun, S. (2016). HMM-DM: identifying differentially methylated regions using a hidden Markov model. Stat. Appl. Genet. Mol. Biol. 15. 10.1515/sagmb-2015-0077.

43. Sun, S., and Yu, X. (2016). HMM-Fisher: identifying differential methylation using a hidden Markov model and Fisher’s exact test. Stat. Appl. Genet. Mol. Biol. 15. 10.1515/sagmb-2015-0076.

44. Saito, Y., Tsuji, J., and Mituyama, T. (2014). Bisulfighter: accurate detection of methylated cytosines and differentially methylated regions. Nucleic Acids Res. 42, e45–e45. 10.1093/nar/gkt1373.

45. Sin, Z., Kinnear, E., Doshi, R., Chatterjee, S., Derbel, H., Guha, P., and Liu, Q. (2025). IPMK depletion influences genome-wide DNA methylation. Biochem. Biophys. Res. Commun. 766, 151874. 10.1016/j.bbrc.2025.151874.

46. Li, X., Liu, Y., Salz, T., Hansen, K.D., and Feinberg, A. (2016). Whole-genome analysis of the methylome and hydroxymethylome in normal and malignant lung and liver. Genome Res. 26, 1730– 1741. 10.1101/gr.211854.116.

47. Loyfer, N., Magenheim, J., Peretz, A., Cann, G., Bredno, J., Klochendler, A., Fox-Fisher, I., Shabi-Porat, S., Hecht, M., Pelet, T., et al. (2023). A DNA methylation atlas of normal human cell types. Nature 613, 355–364. 10.1038/s41586-022-05580-6.

48. Cleveland, W.S. (1979). Robust Locally Weighted Regression and Smoothing Scatterplots. J. Am. Stat. Assoc.

49. Edwards, J.R., O’Donnell, A.H., Rollins, R.A., Peckham, H.E., Lee, C., Milekic, M.H., Chanrion, B., Fu, Y., Su, T., Hibshoosh, H., et al. (2010). Chromatin and sequence features that define the fine and gross structure of genomic methylation patterns. Genome Res. 20, 972–980. 10.1101/gr.101535.109.

50. Ritchie, M.E., Phipson, B., Wu, D., Hu, Y., Law, C.W., Shi, W., and Smyth, G.K. (2015). limma powers differential expression analyses for RNA-sequencing and microarray studies. Nucleic Acids Res. 43, e47–e47. 10.1093/nar/gkv007.

51. Benjamini, Y., and Hochberg, Y. (1995). Controlling the False Discovery Rate: A Practical and Powerful Approach to Multiple Testing. J. R. Stat. Soc. Ser. B Stat. Methodol. 57, 289–300. 10.1111/j.2517-6161.1995.tb02031.x.

52. Guo, S., Diep, D., Plongthongkum, N., Fung, H.-L., Zhang, K., and Zhang, K. (2017). Identification of methylation haplotype blocks aids in deconvolution of heterogeneous tissue samples and tumor tissue-of-origin mapping from plasma DNA. Nat. Genet. 49, 635–642. 10.1038/ng.3805.

53. Affinito, O., Palumbo, D., Fierro, A., Cuomo, M., De Riso, G., Monticelli, A., Miele, G., Chiariotti, L., and Cocozza, S. (2020). Nucleotide distance influences co-methylation between nearby CpG sites. Genomics 112, 144–150. 10.1016/j.ygeno.2019.05.007.

54. Li, Y., Zhu, J., Tian, G., Li, N., Li, Q., Ye, M., Zheng, H., Yu, J., Wu, H., Sun, J., et al. (2010). The DNA Methylome of Human Peripheral Blood Mononuclear Cells. PLOS Biol. 8, e1000533. 10.1371/journal.pbio.1000533.

55. Eckhardt, F., Lewin, J., Cortese, R., Rakyan, V.K., Attwood, J., Burger, M., Burton, J., Cox, T.V., Davies, R., Down, T.A., et al. (2006). DNA methylation profiling of human chromosomes 6, 20 and 22. Nat. Genet. 38, 1378–1385. 10.1038/ng1909.

56. Abascal, F., Acosta, R., Addleman, N.J., Adrian, J., Afzal, V., Ai, R., Aken, B., Akiyama, J.A., Jammal, O.A., Amrhein, H., et al. (2020). Expanded encyclopaedias of DNA elements in the human and mouse genomes. Nature 583, 699–710. 10.1038/s41586-020-2493-4.

57. Perez, G., Barber, G.P., Benet-Pages, A., Casper, J., Clawson, H., Diekhans, M., Fischer, C., Gonzalez, J.N., Hinrichs, A.S., Lee, C.M., et al. (2025). The UCSC Genome Browser database: 2025 update. Nucleic Acids Res. 53, D1243–D1249. 10.1093/nar/gkae974.

58. Akalin, A., Franke, V., Vlahoviček, K., Mason, C.E., and Schübeler, D. (2015). genomation: a toolkit to summarize, annotate and visualize genomic intervals. Bioinformatics 31, 1127–1129. 10.1093/bioinformatics/btu775.

59. Regulatory Feature: ENSR1_BPMG https://useast.ensembl.org/Homo_sapiens/Regulation/Summary?db=core;fdb=funcgen;r=1:3664875-3667889;rf=ENSR1_BPMG.

